# Optogenetic recruitment of hypothalamic corticotrophin-releasing-hormone (CRH) neurons reduces motivational drive

**DOI:** 10.1101/2023.02.03.527084

**Authors:** Caitlin S Mitchell, Erin J Campbell, Simon D Fisher, Laura M Stanton, Nicholas J Burton, Amy J Pearl, Gavan P McNally, Jaideep S Bains, Tamás Füzesi, Brett A Graham, Elizabeth E Manning, Christopher V Dayas

## Abstract

Impaired motivational drive is a key feature of depression. Chronic stress is a known antecedent to the development of depression in humans and depressive-like states in animals. Whilst there is a clear relationship between stress and motivational drive, the mechanisms underpinning this association remain unclear. One hypothesis is that the endocrine system, via corticotropin-releasing hormone (CRH) in the paraventricular nucleus of the hypothalamus (PVN; PVN^CRH^), initiates a hormonal cascade resulting in glucocorticoid release, and that excessive glucocorticoids change brain circuit function to produce depression-related symptoms. Another, mostly unexplored hypothesis is that the direct activity of PVN^CRH^ neurons and their input to other stress- and reward-related brain regions drives these behaviours. To further understand the direct involvement of PVN^CRH^ neurons in motivation, we used optogenetic stimulation to activate these neurons one hour/day for 5 consecutive days and showed increased acute stress-related behaviours and long-lasting deficits in the motivational drive for sucrose. This was associated with increased Fos-protein expression in the lateral hypothalamus (LH). Direct stimulation of the PVN^CRH^ inputs in the LH produced a similar pattern of effects on sucrose motivation. Together, these data suggest that PVN^CRH^ neuronal activity may be directly responsible for changes in motivational drive and that these behavioural changes may, in part, be driven by PVN^CRH^ synaptic projections to the LH.

## Introduction

Mood disorders, including depression, represent a major health and economic burden (1). Depression has a complex aetiology and encompasses a variety of symptoms such as anhedonia, despair, and reduced motivation. Unfortunately, these symptoms often impede the pursuit of and participation in treatment for depression (2). Chronic stress is a key contributor to the onset of depression in humans and can provoke depressive-like behaviour in rodents (3–5). Indeed, hyperactivation of the neuroendocrine arm of the stress response, the hypothalamic-pituitary-adrenal (HPA) axis, has been reported in approximately 50% of individuals with depression (6). Further, positive clinical outcomes after treatment with antidepressants has been associated with the normalization of stress hormone levels (7, 8). Similar results have been reported in rodents, with repeated exposure to low doses of corticosterone in rats resulting in a depressive-like behavioural phenotype which was reversed by chronic antidepressant treatment (9, 10).

While there is a strong relationship between impaired motivational drive and stress-related neuroendocrine activation, much of the evidence linking neuroendocrine activation with distinct behavioural changes is correlational (7–9, 11, 12). Corticotropin-releasing hormone (CRH) neurons in the paraventricular nucleus of the hypothalamus (PVN) constitute the apex of the HPA axis, initiating a hormonal cascade (13) that culminates in glucocorticoid secretion. In rodents, deficits in preference for sucrose as well as other indices, such as reduced sociability, can be replicated by artificially raising glucocorticoids over several weeks to stress-like levels (9, 14), thus implicating glucocorticoids in the manifestation of these depressive-like behaviors. Whilst glucocorticoids have been functionally implicated in depressive-like behaviours, direct evidence of a necessary rather than an associative role for activation of PVN^CRH^ neurons in these behaviours, particularly motivation, is lacking. An alternative hypothesis that has recently gained support is that such stress-induced behaviours do not require endocrine signalling, and that PVN^CRH^ neurons also *directly drive* altered motivational states through synaptic rather than endocrine actions (15). Interestingly, emotional and physiological stressors robustly activate PVN^CRH^ neurons (16, 17), whereas real-time recording of these neurons using fibre photometry has shown that these cells are inhibited by positive rewards such as sucrose (18). These data suggest that stress and motivation interact to determine the activity of these neurons. Further, a recent study that directly manipulated PVN^CRH^ neurons using optogenetics showed that acute stress-related behaviours require an excitatory, glutamatergic projection to the lateral hypothalamus (LH) that occurs on a timescale consistent with synaptic rather than endocrine events (19). In parallel work, we have demonstrated that the LH is an important substrate of chronic stress-induced motivational disturbances (20). Together these studies suggest a more nuanced role for PVN^CRH^ neurons in motivated behaviour, potentially via their projections to brain regions critical for motivation such as the LH or ventral tegmental area (VTA) (21).

Accordingly, here we aimed to test whether directly activating PVN^CRH^ neurons can induce long-lasting motivational disturbances, as assessed by sucrose self-administration, under low and high effort reinforcement conditions in CRH-Cre transgenic mice. We found that repeated optogenetic activation of PVN^CRH^ neurons increased acute stress-like behaviours but produced a long-lasting reduction in motivation for sucrose. This pattern of effects was also observed following direct stimulation of the PVN^CRH^ terminals located in the LH. Together our data provide the first direct demonstration that repeated experimental activation of PVN^CRH^ neurons is sufficient to produce impairments in motivation and suggest that these effects are likely underpinned by PVN^CRH^ neuron projections to brain centers involved in motivation as well as HPA endocrine control.

## Methods

### Ethics statement

All procedures were performed in accordance with the Prevention of Cruelty to Animals Act (2004), under the guidelines of the National Health and Medical Research Council (NHMRC) Australian Code of Practice for the Care and Use of Animals for Experimental Purposes (2013). Animal ethics were approved by The University of Newcastle Animal Ethics Committee.

### Animals

B6(Cg)-CRH^tm1(cre)Zjh^/J (CRH-IRES-Cre) and B6.Cg-Gt(ROSA)26Sor^tm14(CAG-TdTomato)Hze^/J (Ai14; tdTomato) were obtained from Jackson Laboratory (stock numbers 012704 and 007914 respectively). Pairs of homozygous CRH-IRES-Cre and tdTomato mice were crossed to produce heterozygous CRH-IRES-Cre;tdTomato (CRH-Cre::tdTomato) mice. The selective expression of tdTomato in PVN^CRH^ neurons has been previously characterized by our laboratory, consistent with other work (22–24). Our preliminary work showed that the electrophysiological profile and expression of CRH neurons in the PVN and amygdala to be identical to previous studies (25). A total of 40 CRH-Cre and CRH-Cre::tdtomato mice were used in all experiments (8-13 weeks old at beginning of experimentation; 24 female and 16 male). In Experiment 1, 5 animals were excluded from the final analysis due to incorrect fibre optic probe targeting. In Experiment 2, stringent exclusion criteria were set for PVN virus and LH probe placements. Accordingly, the number of animals excluded due to incorrect targeting was 15 due to the complexity of bilateral targeting of the LH and PVN viral hit rate. Food and water were available *ad libitum* and all mice were maintained on a reverse 12-hour light/dark cycle (0700 lights off).

### Experiment 1: Effect of optogenetic recruitment of PVN^CRH^ neurons on motivation for sucrose, Fos-protein activity and stress-related behaviours

Refer to **Figures 1A-C** for the experimental procedures for Experiment 1. Precise details can be found under the subheadings below. Briefly, mice received either YFP control (n=13) or ChR2 (n=15) viral injection surgery targeting the PVN. Fibre optic probe placement occurred during the same surgery in 8 YFP control mice and 9 ChR2 mice (Cohort 2). This was followed by operant training for sucrose under a fixed ratio 1 (FR1) schedule of reinforcement for 10 days then FR3 training for 8 days. Fibre optic probe surgery followed operant training in 5 YFP and 6 ChR2 mice (Cohort 1). All mice were allowed at least 7 days to recover from surgery. Following this, mice were exposed to 2 days of FR3 training and 3 days of progressive ratio (PR) training (termed pre-stimulation training). This was followed by 5 days of chronic PVN^CRH^ optogenetic stimulation consisting of 1hour/day, 10Hz, 10ms pulse width, 15mW, 30 seconds on 30 seconds off. During optogenetic stimulation, stress-related behaviours were recorded in each trial for the first 10 minutes. Following stimulation, mice were split into 2 cohorts, both exposed to 2 FR3 sessions followed by either 3 or 7 PR sessions. Following PR sessions, mice were exposed to a final PVN^CRH^ optogenetic stimulation session and perfused 2 hours later for Fos-protein immunohistochemistry.

**Figure 1.**
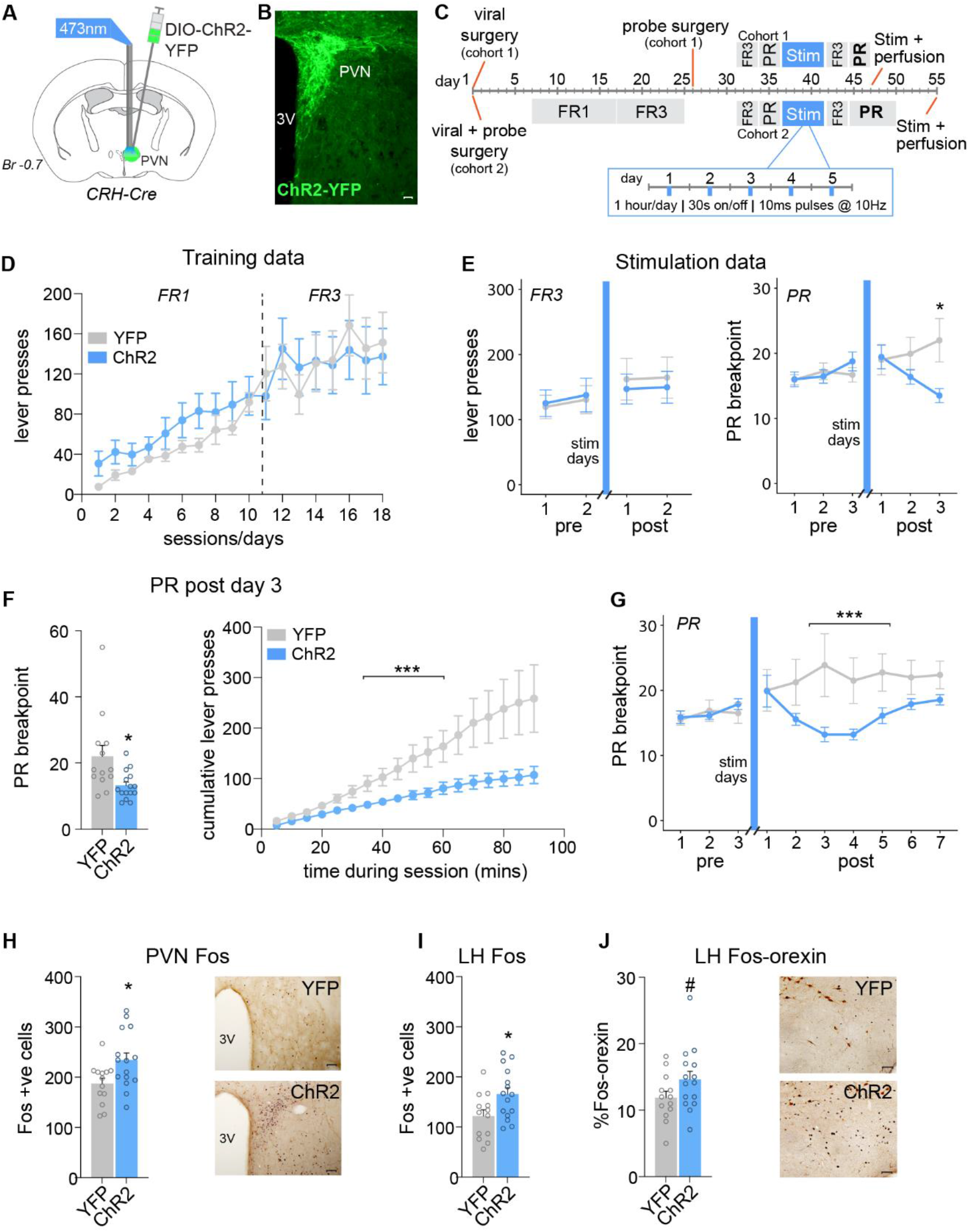
Repeated optogenetic activation of PVN^CRH^ neurons reduces the motivation for sucrose. **A**, CRH-Cre transgenic mice were injected with DIO-ChR2-YFP or the YFP control virus, and implanted with a fibre optic probe, above the PVN. **B**, ChR2-YFP expression was observed in the PVN. Scale bar, 50μm. **C**, Experimental design for Experiment 1 including timing of operant training, surgery and optogenetic stimulation. **D**, There was no difference between mice injected with YFP control virus versus ChR2 virus in the number of active lever presses across FR1 and FR3 training days. **E**, There was no effect of repeated PVN^CRH^ photostimulation on the number of active lever presses under a FR3 schedule of reinforcement. Following repeated PVN^CRH^ photostimulation, there was a reduction in PR breakpoint in ChR2 mice compared to controls. *p<0.05. **F**, ChR2 mice had a significant reduction in PR breakpoint on Day 3 post photostimulation, compared to YFP controls. *p<0.05. This effect was also apparent when lever press data was broken down into 5min time bins. ***p<0.001 Time x Virus interaction. **G**, The reduction in PR breakpoint following PVN^CRH^ photostimulation in ChR2 mice persisted for up to 7 days post stimulation, although the effect weakened over time. ***p<0.001 Day x Virus interaction effect. **H**, There was a significant increase in the number of Fos-positive cells in the PVN in ChR2 mice compared to YFP controls following PVN^CRH^ stimulation. *p<0.05. Scale bar, 200μm. **I**, There was a significant increase in the number of Fos-positive cells in the LH in ChR2 mice compared to YFP controls following PVN^CRH^ stimulation. *p<0.05. **J**, There was a trend towards a significant increase in the percentage of Fos-positive neurons that were orexin positive in the LH in ChR2 mice compared to controls following PVN^CRH^ photostimulation. #p=0.098. Scale bar, 50μm. PVN, paraventricular nucleus of the hypothalamus; LH, lateral hypothalamus; PR, progressive ratio; FR, fixed ratio; ChR2, channelrhodopsin-2; YFP, yellow fluorescent protein; stim, optogenetic stimulation of PVN^CRH^ neurons; Br, bregma.

### Experiment 2: Effect of optogenetic recruitment of the PVN^CRH^ to LH pathway on motivation for sucrose and stress-related behaviours

Refer to **Figures 3A-B** for the behavioural procedure for Experiment 2. Precise details of procedures are described below. Briefly, mice received either YFP control (n=6) or ChR2 viral injection surgery (n=6) targeting the PVN. In the same surgery, mice were implanted with fibre optic probes targeting the LH. This was followed by 7 days recovery and 10 days FR1 and 8 days FR3 training for sucrose. Mice were given a 7-day break from operant training to ensure the same timeline as Experiment 1. This was followed by 2 days of FR3 pre-stimulation training and 3 days of PR prestimulation training. Chronic PVN terminal optogenetic stimulation in the LH consisted of 5 days at 20Hz, 10ms pulse width, 15mW, 30 seconds on 30 seconds off for 1 hour/day. Stress-related behaviours were monitored during stimulation sessions for the first 10 minutes. Following stimulation, mice were exposed to 2 FR3 operant sessions and 3 PR sessions.

### Surgery

CRH-Cre or CRH-Cre::tdTomato mice were anesthetized with isoflurane (5% induction, 2% maintenance) before being placed into a stereotaxic frame (Stoelting Instruments). Mice received stereotaxic PVN-directed injections of either AAV5-DIO-ChR2-YFP or AAV5-DIO-YFP (Addgene; anteroposterior (AP): −0.7, mediolateral (ML): ± 0.25, dorsoventral (DV): −4.8) to selectively transduce CRH neurons (**Figures 1A, 1B**). 0.2μl was injected at a rate of 0.15μl/min using a 2μl Hamilton Neuros syringe (30G) attached to a Stoelting Quintessential Stereotaxic Injector pump. Craniotomies were closed with BoneWax (Ethicon, NJ, USA) and the scalp incision secured with 3M Vetbond and staples (AutoClip, Fine Science Tools, VIC, AUS) for Experiment 1 cohort 1. For placement of fibre optic probes, two craniotomies were drilled and stainless-steel screws inserted (Avivamann Optical Group, WA, AUS). Probe and screws were secured to the skull with 3M Vetbond and dental glue (Dentsply Sirona Pty Ltd, NSW, AUS). Mono fibre optic cannula placement (6 mm; 400 μm core; Slim Magnetic Receptor (SMR); 0.37 NA; Doric Lenses, QC, CAN) was either dorsal to the PVN in Experiment 1 (AP: −0.7, ML: 0.0, DV: −4.3) or directly in the LH in Experiment 2 during the viral surgery (bilateral: AP: −1.2, ML: +/−1.0, DV: −4.75) (**Figures 1A, 3A**). After surgery, mice were given a subcutaneous injection of Ketoprofen (5ml/kg) for analgesia.

### Sucrose self-administration

Mice were first trained to self-administer sucrose (10% w/v) in daily 30-minute operant conditioning sessions under a fixed ratio 1 (FR1) schedule of reinforcement (Med Associates, VT, USA). A response on the active (right) lever resulted in the delivery of 10μl of sucrose into a receptacle followed by a 4 second light cue located above the lever to indicate sucrose availability. Following 10 days of FR1 training, mice were moved to a FR3 schedule of reinforcement for the next 8 days where an inactive (left) lever was introduced which had no scheduled consequence (**Figure 1C**). After FR3 training, mice had 7 days rest from operant self-administration to allow for recovery from probe surgery from Experiment 1 cohort 1. This was followed by 2 additional FR3 operant sessions and 3 PR sessions. PR sessions occurred daily for 90 minutes using a reinforcement schedule where the number of lever presses required for sucrose reward delivery increased by 1 following each reward i.e. 1,2,3,4 …. Etc. (26, 27). PR breakpoint was defined as the highest response ratio completed. Both FR3 and PR testing was undertaken before and after repeated optogenetic stimulation (**Figures 1C, 3B**). All behavioural testing took place between 8am and 6pm daily.

### *In vivo* optogenetics

Optogenetic experiments aimed to examine the behavioural activity and Fos-protein expression produced by the optogenetic activation of PVN^CRH^ neurons or PVN^CRH^ terminals in the LH (28). One week prior to behavioural testing, animals were handled twice daily and habituated to their testing apparatus (Perspex arena; 42.2 cm L x 72 cm W x 60 cm H, with clean corn cob floor material). On the test day, animals were attached to fibre optic patch cords (400 μm core; FCM-SMC; 0.37 numerical aperture (NA); Doric Lenses) via their implanted mono fibre optic cannula. The opposing end of the patch cord was connected to a FC-FC fibre optic 1 x 1 rotary joint (Doric Lenses). A 473 nm DPSS laser (Laserglow Technologies, ON, CAN) was used to deliver light into the rotary joint via a patch cord with FC/FC connections on each end. Laser parameters were controlled using a Master-8 pulse stimulator (A.M.P.I, JRS, IL).

Power output from the end of FCM-SMC patch cord (15mW) was determined using a photodiode power sensor connected to a console (Thorlabs, NJ, USA). During each session, mice were given 30 minutes following patch cable attachment before the laser was turned on for 60 minutes.

### Acute behavioural analysis

For all animals, behaviour was recorded for 12 minutes commencing 2 minutes before blue light stimulation (2 minutes baseline + 10-minute laser stimulation analysis). Behavioural data was analysed using a video annotator program (developed by Dr Toni-Lee Sterley – Hotchkiss Brain Institute, CAN) and used to quantify the duration of six distinct behaviours (digging, grooming, rearing, walking, jumping, inactive) (19). Behavioural video analysis was undertaken by two individuals blind to treatment condition. The output from this program generated a record of all behaviours displayed over the 10-minute period in Microsoft Excel.

### Immunohistochemistry and microscopy

Following the completion of the experiments, mice were euthanised with sodium pentobarbitone (0.2ml intraperitoneal, I.P, Virbac, AUS) 2 hours after the commencement of their final 1-hour optogenetic stimulation session, and perfused with 20ml of 0.1M Phosphate Buffer Solution (PBS) and 50ml of 4% paraformaldehyde (PFA). Brains were extracted and postfixed in 4% PFA at 4°C for 2 hrs, then transferred to 30% sucrose in 0.1M PBS for 48 hrs at 4°C. Brains were frozen on dry ice and stored at −80°C until sectioning. Coronal sections 40 μm thick were cut on a cryostat (Leica Biosystems CM1900) in a 1 in 4 series and stored in a 0.1 M PBS solution containing 0.1% sodium azide at 4°C. For the detection Fos-protein in Experiment 1, tissue sections were incubated in a solution of PBS containing 1 % Triton X-100, 2% normal horse serum and primary antibody (1:8000, Phospho-c-Fos, rabbit monoclonal; Cell Signaling Technology) for 48 hours at 4 °C. Sections were then washed 3 times for 10 minutes in PBS before incubation with the secondary antibody (1:1000, donkey anti-rabbit IgG, Jackson ImmunoResearch, PA, USA) for 2 hours at room temperature. After washing to remove unbound secondary antibody, sections were incubated for 1 hour using a Vectastain ABC kit (Vector Laboratories, CA, USA) followed by incubation with diaminobenzodine in 2% filtered nickel sulfate for 15 minutes. Visualisation of Fos-protein was achieved by the addition of glucose oxidase.

This tissue was then co-labelled for orexin-A. To co-label without cross-reactivity from rabbit polyclonal antibodies, an Avidin/Biotin Blocking kit was used (Vector Laboratories, CA, USA) before incubating tissue in orexin primary antibody (1:10000, rabbit monoclonal, Phoenix Pharmaceuticals, CA, USA) for 48 hours at 4 °C. The procedures and reagents for visualization of cytoplasmic orexin immunoreactivity, ABC kit and visualization steps are the same as for Fos-protein with the exception that 2% filtered nickel sulfate was replaced with 0.1% acetate buffer. Following completion of immunohistochemical procedures, brain sections were mounted on chrome alum-treated slides, dehydrated in a series of ethanol and xylene solutions and coverslipped.

Photomicrographs of brain sections were made using Olympus CellSens Software (version 1.3) on an Olympus BX51 microscope at 10x objective. Bilateral cell counting was undertaken using iVision (Biovision) computer program for PVN slices (Bregma − 0.6mm to −0.8mm). Image J was used to quantify Fos-positive and orexin-positive cells in LH sections (Bregma −1.22 to −1.7). Coronal brain sections were chosen according to Paxinos and Franklin’s Mouse Brain in Stereotaxic Coordinates 6^th^ edition (29) and quantification was undertaken by one observer, blind to treatment.

PVN and LH sections from a second series of brain tissue was mounted onto microscope slides and coverslipped using a 0.1M PBS and glycerol solution. Probe placement and viral expression were then visualized using an Olympus BX51 microscope. Please refer to **Figure S1** for fibre optic probe placements.

### Statistical analysis

Statistical analyses were conducted using JASP V16.2. Sucrose lever pressing was analysed using an analysis of variance (ANOVA) with the between-subjects factor of Virus (YFP, ChR2) and the within-subjects factor of Day for initial FR1 and FR3 operant training, and two within subject factors, day and stimulation period (pre vs post) for FR3 and PR operant testing, as well as 5min time bins for PR responding on Day 3. A one-way between-subjects ANOVA was used to assess the effect of optogenetic PVN^CRH^ neuron recruitment on Fos-protein activity. Stress-related behaviours were analysed using ANOVA with the between-subjects factor of Virus (YFP, ChR2) and the within-subjects factor of Day. Tukey post-hoc comparisons were used to differentiate any significant interactions.

## Results

### Effect of optogenetic recruitment of PVN^CRH^ neurons on motivation for sucrose

CRH-Cre transgenic mice were injected with a Cre-dependent excitatory opsin (ChR2) into the PVN (**Figures 1A-B**). Additionally, a fibre optic probe was implanted immediately dorsal to the PVN to allow optogenetic activation of CRH neurons in the PVN (**Figure 1C**). Mice were initially trained to lever press for sucrose by increasing fixed ratio schedules (**Figure 1C**). Analysis of active lever presses during operant training revealed no significant interaction of Virus and Day during the FR1 schedule (F_9,234_ = 0.305, p = 0.973) or FR3 schedule of reinforcement (**Figure 1D;** F_7,182_ = 0.731, p = 0.646). Following operant training, mice were assessed for their motivation for sucrose under both low effort (FR3) and high effort (PR) schedules of reinforcement at baseline and following 5 days of photostimulation of PVN^CRH^ neurons (**Figure 1C;** “Stim” sessions). Under FR3 conditions, there was no effect of repeated PVN^CRH^ photostimulation on the number of active lever presses for sucrose (**Figure 1E;** F_1,26_ = 0.315, p = 0.865). There was no effect of Virus on inactive lever presses during FR3 training days or PR sessions (**Figure S2**). However, under higher effort PR conditions, there was a significant effect of repeated PVN^CRH^ photostimulation on the number of active lever presses for sucrose (F_2,52_ = 14.06, p < 0.0001). There was also an effect of repeated PVN^CRH^ stimulation on breakpoint during PR sessions (**Figure 1E;** F_2,52_ = 14.20, p < 0.0001). Tukey post-hoc comparisons revealed significant differences in breakpoint between YFP control animals and ChR2 animals on Day 3 post-stimulation (**Figure 1F;** p = 0.025). Closer inspection of 5min time bin active lever press data on Day 3 post-stimulation revealed a significant interaction of Virus and Time bin on active lever presses (**Figure 1F;** F_17,442_ = 5.105, p < 0.001). Interestingly, when the protocol was extended to 7 days of PR, in a second cohort of mice, the effect of repeated PVN^CRH^ stimulation on breakpoint persisted, but began to return to control levels by Day 7 (**Figure 1G;** F_9,135_ = 3.379, p < 0.001).

### Effect of optogenetic recruitment of the PVN^CRH^ neurons on Fos-protein activity

On the day after the final PR session, mice underwent a final photostimulation session and were perfused 2 hours later. Immunohistochemical analysis revealed an increased number Fos-positive cells in the PVN (F_1,26_ = 6.011, p = 0.021) and LH (F_1,26_ = 5.778, p = 0.024) in ChR2 mice compared to YFP controls (**Figures 1H; 1I**). There was a trend for an effect of Virus on the percentage of Fos-positive orexin cells in the LH (**Figure 1J;** F_1,26_ = 2.942, p = 0.098).

### Effect of optogenetic recruitment of the PVN^CRH^ neurons on stress-related behaviours

We determined the effect of PVN^CRH^ photostimulation on stress-like behaviours across the five days during the first 10 minutes of 473 nm blue light exposure. Behavioural analysis during the photostimulation sessions demonstrated a significant effect of Virus and Behaviour on the time spent engaging in each stress-related behaviour (**Figures 2A-G;** F_6,189_ = 18.260, p < 0.0001). Tukey post-hoc comparisons showed that ChR2 mice exhibited significantly increased digging and grooming behaviours than YFP controls across the five sessions (p’s < 0.05). Additionally, ChR2 mice exhibited reduced rearing and exploratory behaviours (walking) compared to YFP controls (p’s < 0.05).

**Figure 2.**
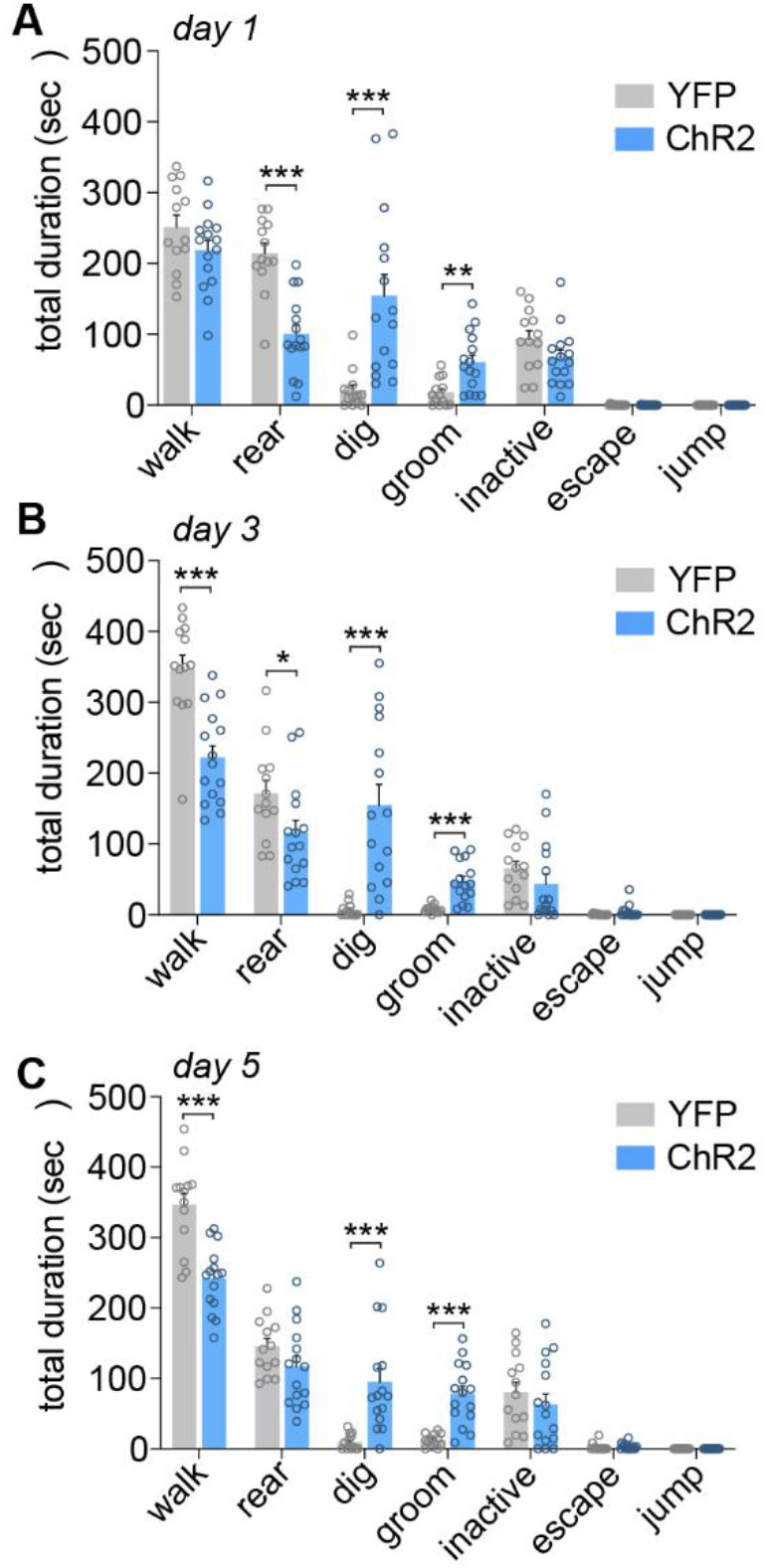
Optogenetic stimulation of PVN^CRH^ neurons increases stress-related behaviours. **A**, During the first PVN^CRH^ photostimulation session, a shift in stress-related behaviours was observed in ChR2-mice compared to YFP controls including reduced rearing behaviour, increased digging behaviour and increased grooming. **p<0.01, ***p<0.001 post-hoc analyses. **B**, During the third photostimulation session, ChR2 mice showed increased stress-related behaviours compared to YFP controls including reduced rearing behaviour, increased digging behaviour, increased grooming and reduced walking. *p<0.05, ***p<0.001 post-hoc analyses. **C**, During the fifth PVN^CRH^ photostimulation session, ChR2 mice showed increased stress-related behaviours compared to YFP controls including increased digging behaviour, reduced walking, and increased grooming. ***p<0.001 post-hoc analyses. PVN, paraventricular nucleus of the hypothalamus.

### Effect of optogenetic recruitment of the PVN^CRH^ to LH pathway on motivation for sucrose

Previous work has shown that the LH is a target of PVN^CRH^ terminals, and that activation of this pathway can elicit acute stress-related behaviours (19). Further, in experiment 1 optogenetic photostimulation of PVN^CRH^ cell bodies resulted in increased LH Fos. Thus, we next asked whether repeated stimulation of this circuit would suppress the motivation for sucrose in a similar manner to cell body stimulation. CRH-Cre transgenic mice were injected with DIO-ChR2 into the PVN, and fibre optic probes were implanted bilaterally immediately dorsal to the LH (**Figure 3A-B**). Mice completed the same sucrose training and photostimulation sessions as conducted in Experiment 1 (**Figure 3B**), but instead, targeting PVN^CRH^ terminals in the LH. Analysis of active lever presses during operant training showed no significant effect of Virus and Day during the FR1 schedule (F_9,90_ = 0.713, p = 0.696) or FR3 schedule of reinforcement (**Figure 3C;** F_7,70_ = 0.183, p = 0.988). Following operant training, mice were again assessed for their motivation for sucrose under both low effort (FR3) and high effort (PR) schedules of reinforcement at baseline and following 5 days of photostimulation of PVN^CRH^ terminals in the LH (**Figure 3B;** “Stim” sessions). Under low effort, FR3 conditions, there was no effect of repeated PVN^CRH^ terminal photostimulation in the LH on the number of active lever presses for sucrose (**Figure 3D;** F1,10 = 0.071, p = 0.795). There was also no significant effect of repeated PVN^CRH^ terminal stimulation in the LH on the number of active lever presses for sucrose under higher effort PR conditions (F_5,50_ = 1.569, p = 0.186). Following PVN^CRH^ terminal stimulation in the LH, there was a trend for altered breakpoint during PR sessions (**Figure 3D;** F_2,20_ = 3.475, p = 0.051). Separate analysis of post-stimulation breakpoint revealed a significant Day x Virus interaction (F_2,20_ = 4.718, p = 0.021), suggesting a similar pattern of effects to cell body stimulation in Figure 1F. The largest effect of PVN^CRH^ photostimulation observed in Experiment 1 occurred on Day 3 of PR. Thus, we also decided to closely examine this timepoint following optogenetic recruitment of the PVN^CRH^ to LH pathway. There was no significant effect of PVN^CRH^ terminal stimulation in the LH on PR breakpoint when comparing pre-stimulation versus poststimulation on Day 3 of PR testing (**Figure 3E;** F_1,10_ = 2.781, p = 0.126). There was a trend towards a significant reduction in the number of active lever presses for sucrose in ChR2 mice compared to YFP controls on PR Day 3 post-stimulation (**Figure 3F;** F_1,10_ = 3.517, p = 0.090). Time bin active lever press data on Day 3 post-stimulation revealed a significant effect of Virus and Time bin on active lever presses (**Figure 3G;** F_17,170_ = 2.782, p < 0.001).

**Figure 3.**
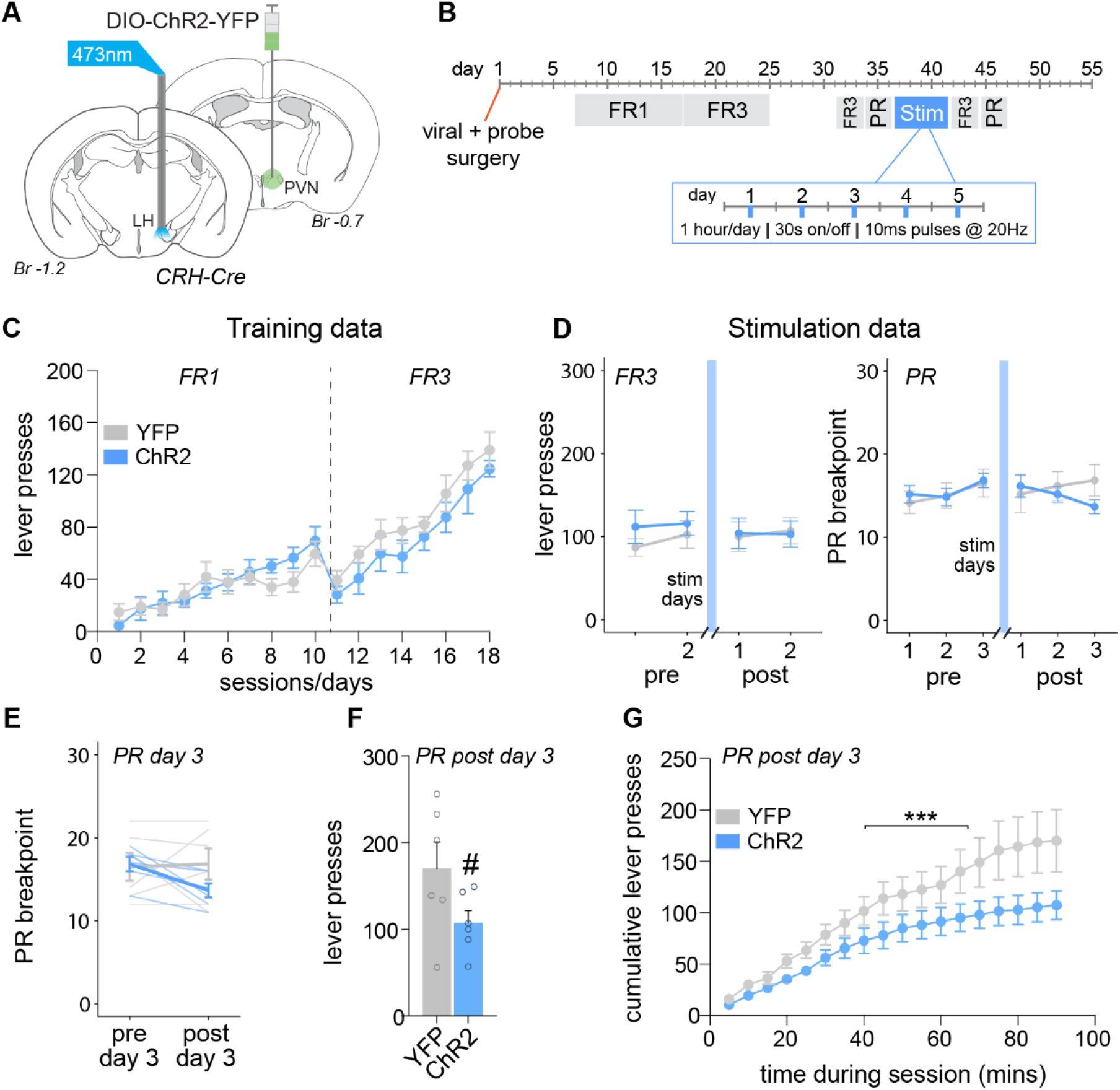
Repeated photostimulation of the PVN^CRH^ to LH pathway modestly reduces the motivation for sucrose. **A**, CRH-Cre transgenic mice were injected with DIO-ChR2-YFP or the YFP control virus into the PVN and implanted with fibre optic probes bilaterally in the LH. **B**, Experimental design for Experiment 2 including timing of operant training, surgery and optogenetic stimulation. **C**, There was no difference between mice injected with YFP control virus versus ChR2 virus in the number of active lever presses across FR1 and FR3 training days. **D**, There was no effect of repeated PVN^CRH^ terminal optogenetic stimulation in the LH on the number of active lever presses under a FR3 schedule of reinforcement. There was also no effect of repeated PVN^CRH^ to LH pathway activation on breakpoint under a PR schedule of reinforcement. **E**, There was no effect of photostimulation day on PR breakpoint between ChR2 mice and YFP controls. **F**, There was a trend towards a significant reduction in the number of PR lever presses in ChR2 mice compared to YFP controls on day 3 post-photostimulation. #p=0.09. **G**, When examining cumulative lever press data on day 3 post-photostimulation for PR, there was a significant reduction in the number of lever presses over time in ChR2 mice compared to YFP controls. ***p<0.001. PVN, paraventricular nucleus of the hypothalamus; LH, lateral hypothalamus; progressive ratio (PR); fixed ratio (FR); optogenetic stimulation of PVN^CRH^ to LH pathway (stim); Br, bregma.

### Effect of optogenetic recruitment of the PVN^CRH^ to LH pathway on stress-related behaviours

Similar to PVN^CRH^ stimulation experiments, optogenetic recruitment of the PVN^CRH^ to LH pathway significantly increased stress-related behaviours compared to YFP controls. There was a significant effect of Virus and Behaviour on the time spent engaging in each stress-related behaviour (**Figures 4A-G;** F_6,70_ = 8.24, p < 0.0001). Tukey post-hoc comparisons showed that ChR2 mice had significantly increased digging during the stimulation on day 1 compared to YFP controls (p < 0.05). Additionally, ChR2 mice exhibited increased grooming across the three sessions analysed (p’s < 0.05).

**Figure 4.**
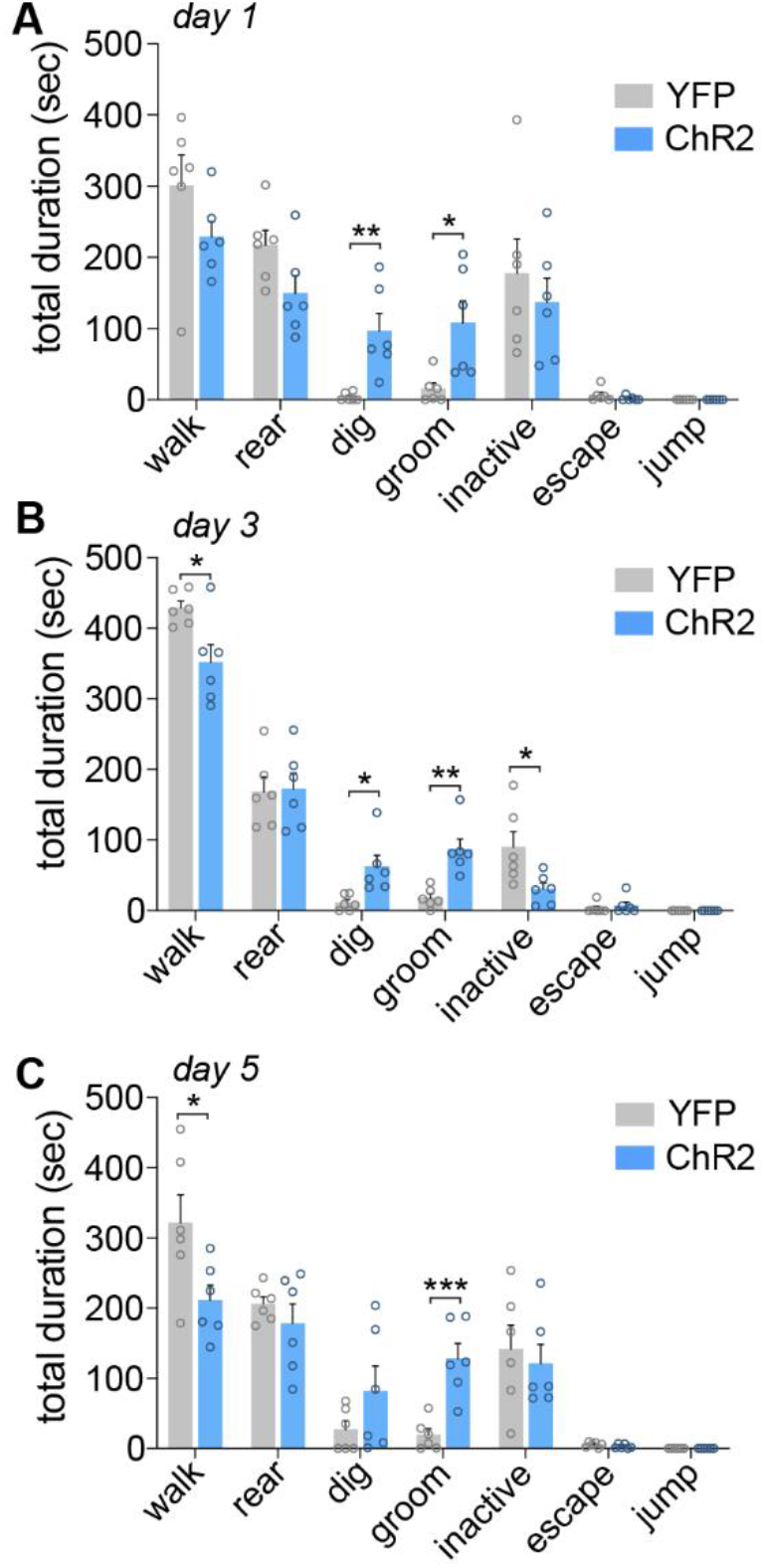
Optogenetic stimulation of the PVN^CRH^ to LH pathway increases stress-related behaviours. **A**, During the first photostimulation session, ChR2 mice showed increased stress-related behaviours compared to YFP controls including increased digging behaviour and increased grooming. *p<0.05; **p<0.01 post-hoc analyses. **B**, During the third PVN^CRH^ to LH pathway photostimulation session, ChR2 mice showed increased stress-related behaviours compared to YFP controls with increased grooming behaviour, increased digging, reduced walking and reduced periods of inactivity. *p<0.05; **p<0.01 post-hoc analyses. **C**, During the fifth photostimulation session, ChR2 mice showed increased grooming behaviour and reduced walking. *p<0.05; ***p<0.001 post-hoc analyses. PVN, paraventricular nucleus of the hypothalamus; LH, lateral hypothalamus.

## Discussion

Stress-induced activation of the HPA axis is strongly associated with mood disorders such as depression, however, a direct link between the activation of PVN^CRH^ neurons that control the HPA axis and impairments in motivational drive has been lacking. Here we show that repeated photoactivation of PVN^CRH^ neurons increased acute stress-related behaviours including grooming and digging, and is sufficient to reduce the motivation to lever press for sucrose, exclusively in a high-effort progressive ratio schedule of reinforcement. Interestingly, this effect persisted for up to 7 days following PVN^CRH^ stimulation. Direct stimulation of PVN^CRH^ neurons increased Fos-protein activity in the PVN and LH, with a trend towards an increase in LH orexin-expressing neurons. Given that recent literature has implicated the PVN^CRH^ to LH pathway in stress-related behaviours and early life stress promotes LH-driven impairments in motivational drive (19, 20), we examined the direct effect of PVN^CRH^ to LH stimulation on the effort to lever press for sucrose. Repeated activation of this pathway produced a similar pattern of changes in PR responding to PVN^CRH^ neuronal cell body stimulation, although not as robustly as activating PVN^CRH^ neuron cell bodies directly. Acutely, photoactivation of PVN^CRH^ terminals in the LH also produced characteristic stress-associated behavioural responses, including increased digging and grooming. Together, these data suggest that PVN^CRH^ neuron activity can precipitate depressive-like behaviours that may extend beyond increased HPA axis activation, as demonstrated by our finding that repeated photostimulation of PVN^CRH^ neurons directly impacts motivational drive.

### PVN^CRH^ activation increased acute stress-related behaviours, reduced motivated responding for sucrose, and altered Fos-protein immunoreactivity

In the current study, we found that repeated activation of PVN^CRH^ neurons selectively suppressed responding for sucrose under a high-effort progressive ratio (PR) schedule of reinforcement, but not under low-effort fixed ratio schedule (FR3). This effect of PVN^CRH^ stimulation on motivational drive persisted for up to 7 days poststimulation, although the effect weakened over time. This finding demonstrates that repeated activation of PVN^CRH^ neurons has a lasting impact on the incentive motivation for sucrose, a phenotype typically observed in people with depression. Whilst we cannot discount the impact of repeated PVN^CRH^ activation on other cognitive domains (particularly those associated with depressed states) (30), an effect of repeated stimulation on FR3 responding was not evident, suggesting relatively selective effects on high effort motivation. Additionally, prior to PVN^CRH^ stimulation, both YFP controls and ChR2 mice showed no differences in FR1 or FR3 operant training, arguing against any potential unexpected behavioural effects of the virus. Interestingly, the impact of stimulation on sucrose responding only began to manifest from day 2 of PR testing. We have observed similar behavioural effects in our previous work assessing the impact of early life stress on the motivation for sucrose (20). In this study, PR breakpoints were similar on day 1 of testing in control and early life stressed groups, however stressed rats showed a significant decrease in motivation from day 2 onwards. This phenotype is consistent with other preclinical work demonstrating that social defeat stress reduces PR responding for saccharin and sucrose consumption up to 5 weeks following stress (31, 32). These preclinical findings, together with our own, align with the human depression literature where chronic stressors have been associated with persistent depressive symptoms, frequent relapse, and poorer prognosis (33, 34). Importantly, our findings suggest a direct role for PVN^CRH^ neurons in these depression-related motivational impairments.

Consistent with recent work (19), we found that optogenetic activation of PVN^CRH^ neurons produced a highly reproducible pattern of stress-related behaviours. We extend this work by demonstrating that these PVN^CRH^ stimulation-evoked behaviours are relatively stable across five days of stimulation. This pattern was characterized by increased grooming and digging, along with reduced walking and rearing behaviour. Grooming is a complex innate behaviour and in the context of stress is thought to be important for de-arousal following stress exposure (35). The increase in digging behaviour we observed during activation of PVN^CRH^ neurons has not been previously reported following a similar experimental design (19). A potential explanation for digging in our experimental preparation is the inclusion of bedding in the test cage, which may have allowed for the expression of this behaviour. Like grooming, digging is an innate rodent behaviour that has potential for pathological repetition and is associated with stressful and anxiogenic situations (36). Digging has conceptual similarity to the anxiety-associated marble burying test, which has been argued to be more appropriately considered an indicative measure of repetitive digging (37, 38). Importantly, behaviours elicited by optogenetic stimulation occurred rapidly, within seconds, therefore these responses are likely driven by synaptic rather than neuroendocrine mechanisms (39). This is supported by our similar acute behavioural observations following selective optogenetic stimulation of PVN^CRH^ terminals in the LH. These findings indicate that PVN^CRH^ neurons mediate fast, stress-relevant behaviours, likely independent of hormonal feedback (19).

Following repeated PVN^CRH^ photostimulation, we observed a significant increase in the number of Fos-positive cells in the PVN and LH. These findings are consistent with work from Fuzesi and colleagues (19), who showed that acute activation of PVN to LH terminals can produce a range of stress-relevant behaviours through direct, excitatory projections to the LH. However, the identity of these LH Fos-positive neurons remains to be determined. Prior evidence from our lab has shown a role of LH orexin in early life stress-induced motivational deficits (20), and in the current study repeated PVN^CRH^ stimulation was associated with a trend towards a significant increase in the percentage of Fos-positive orexin cells. This is interesting as other work using monosynaptic retrograde rabies tracing has shown that CRH directly innervates LH orexin neurons (40). While this effect did not reach statistical significance, there are known limitations with Fos-protein mapping including sensitivity and temporal resolution. Additionally, orexin neurons are spontaneously active and sensitive to many different environmental and behavioural factors, including changes in light, social interaction and energy status (41). Future studies using electrophysiological recordings or Ca^2+^ imaging might be required to detect temporally specific changes in orexin cell activity (19, 20). Finally, we cannot rule out the possibility that other LH neuronal populations may be playing a role including melanin-concentrating hormone neurons and GABAergic neurons (42–44).

### PVN^CRH^ to LH pathway activation modestly reduced motivated responding for sucrose and increased stress-related behaviours

A similar, although weaker, effect on high-effort PR responding was found following repeated photostimulation of PVN^CRH^ terminals in the LH. Photostimulation of PVN^CRH^ terminals in the LH also produced a similar pattern of acute stress-related behaviours, including digging and grooming, as PVN^CRH^ cell body stimulation, consistent with previous literature (19). There is growing interest in the projections of PVN^CRH^ neurons beyond the median eminence, with recent demonstration of important functions outside HPA axis and autonomic activity. Indeed, PVN^CRH^ neurons are known to project to a number of other downstream sites with well described roles in reward and stress-coping behaviours, such as the ventral tegmental area (VTA), locus coeruleus, periaqueductal gray and globus pallidus (45–50). Thus, it may be that activation of additional terminal projections of the CRH neurons is required to produce the full PVN^CRH^ neuronal cell body stimulation effect on motivation (51). In support, acute stress results in CRH release in the VTA, where it has been shown to modulate reward-related behaviour (52). Thus, future research on motivational drive following activation of other reward-related projection pathways of PVN^CRH^ neurons is required.

### Methodological considerations

Our most interesting finding is that direct PVN^CRH^ stimulation produces persistent reductions in motivational drive and an immediate onset of stress-related grooming and digging. This appears to be independent of HPA-axis activity and extends the previous work of Fuzesi and colleagues (19). One limitation of the current study is that we did not assess the involvement of circulating glucocorticoids on motivational drive. Whilst the current experiments do not have a direct read-out of neuroendocrine output, we know CRH neuron activation leads to glucocorticoid secretion, but given the reciprocal connections with the LH we cannot rule out effects of LH terminal stimulation on circulating glucocorticoids (53, 54). Importantly, the key outcome from this work is that activation of CRH neurons in the PVN is sufficient on its own to promote motivational disturbances, suggesting that future studies examining the links between stress and behaviour need to consider both the synaptic and endocrine projections of these cells.

### Conclusions

Here we show ChR2-mediated stimulation of PVN^CRH^ neurons produced a long-lasting reduction in the motivation for a natural reward, distinct from a general behavioural deficit in sucrose operant responding as demonstrated by lack of effects on low effort behaviour. These results are consistent with literature suggesting that repeated activation of stress responses mediated by the PVN can lead to depressive-like behaviours in rodents and humans (55, 56). Overall, these data have important implications for the identification of novel hypothalamic circuits that govern stress-induced changes in mood and motivation. These current advances may aid in developing more effective treatments for debilitating symptoms of neuropsychiatric diseases such as depression, through selective targeting of these PVN projection pathways.

## Acknowledgements

We would like to thank Dr Toni-Lee Sterley for sharing the behaviour annotator she developed for acute behavioural analysis. This work was supported by grants to CVD from the NHMRC Australia

## Supplementary figures

**Figure S1.**
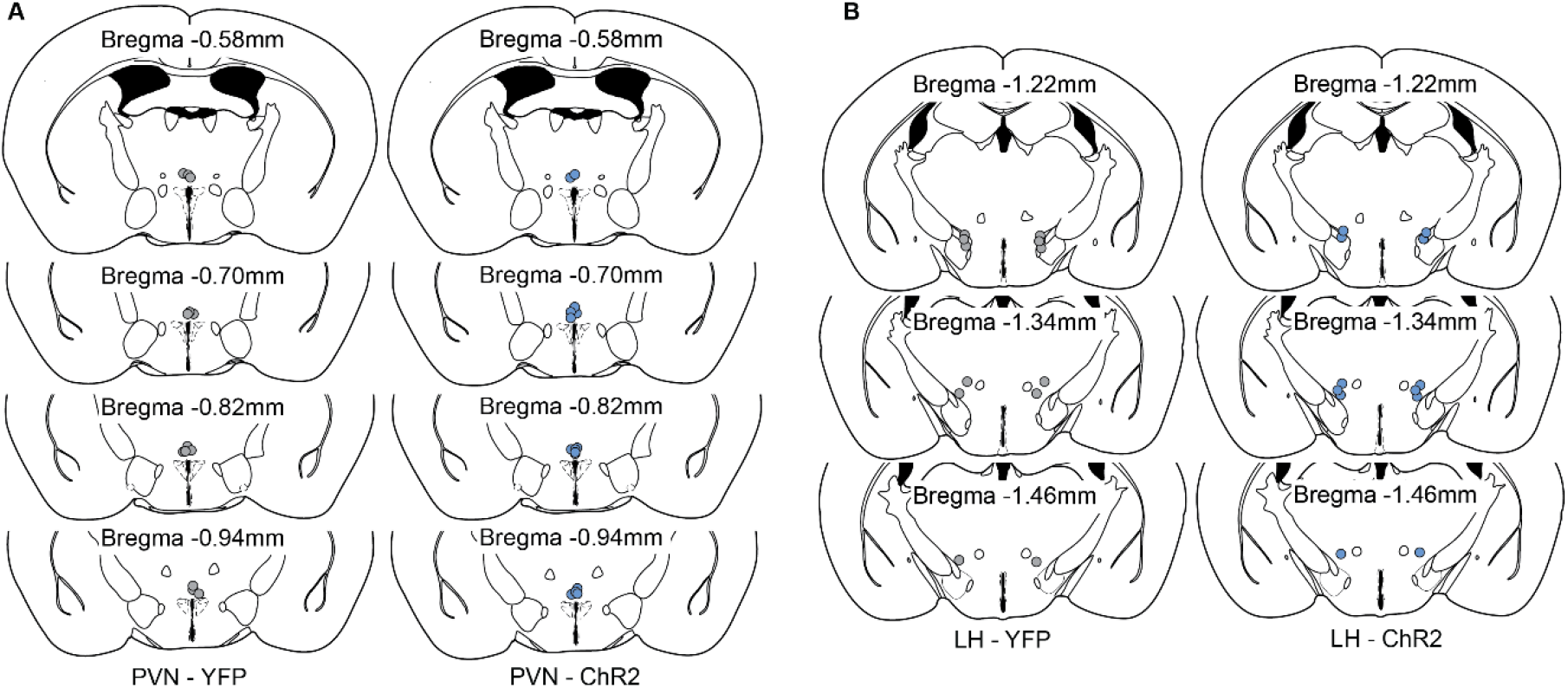
Fibre optic probe placement for PVN^CRH^ stimulation experiments and PVN^CRH^ to LH experiments. **A**, Fibre optic probe placement above the PVN for YFP control mice and ChR2 mice for repeated PVN^CRH^ stimulation experiments. **B**, Fibre optic probe placement relative to Bregma above the LH for YFP control mice and ChR2 mice for repeated PVN^CRH^ to LH stimulation experiments. Mouse brain coordinates were based on Paxinos & Franklin, 2001. PVN, paraventricular nucleus of the hypothalamus; LH, lateral hypothalamus.

**Figure S2.**
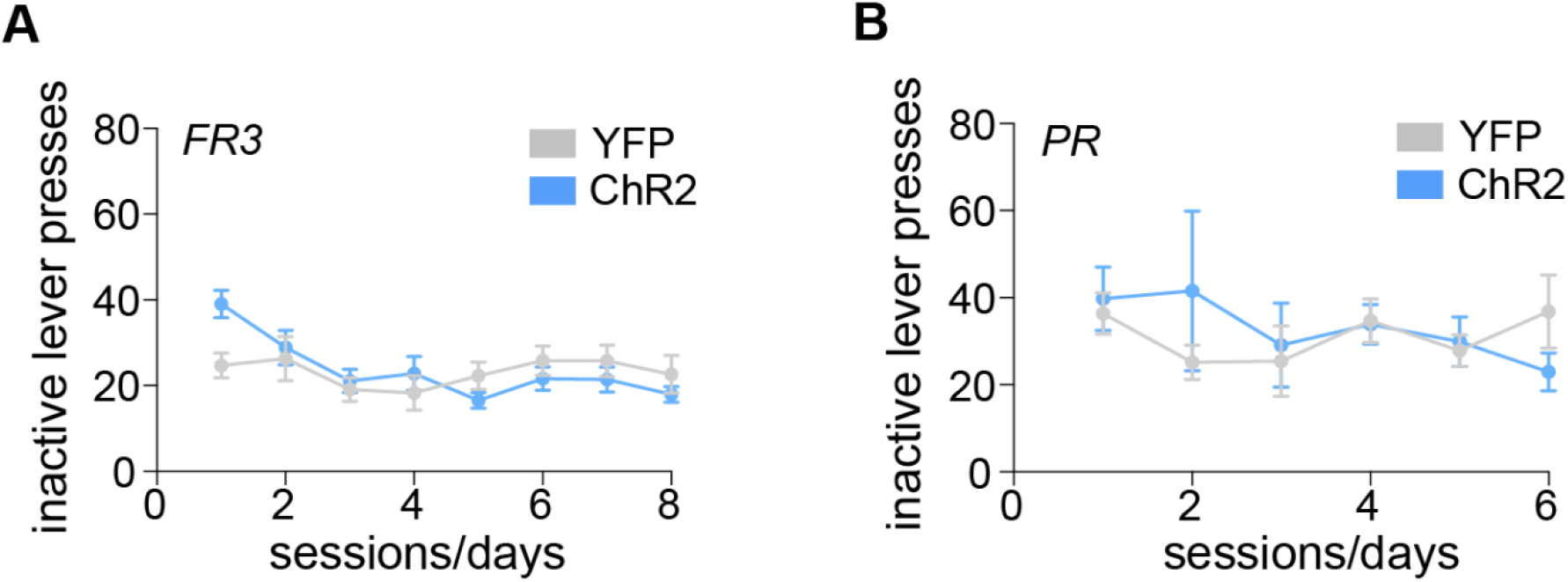
Optogenetic stimulation of PVN^CRH^ neurons does not impact inactive lever pressing during operant training for sucrose. **A**, There was no difference between mice injected with YFP control virus versus ChR2 virus in the number of inactive lever presses across FR3 training days. **B**, There were no differences in the number of inactive lever presses during PR training between YFP control mice and ChR2 mice (sessions 1-3). There was also no effect of repeated PVN^CRH^ photostimulation on the number of inactive lever presses during PR sessions between YFP control mice and ChR2 mice (sessions 4-6). Progressive ratio (PR); fixed ratio (FR).

