## Supplementary figures and images for "Optogenetic recruitment of hypothalamic corticotrophin-releasing-hormone (CRH) neurons reduces motivational drive"

### Supplement Figure 1

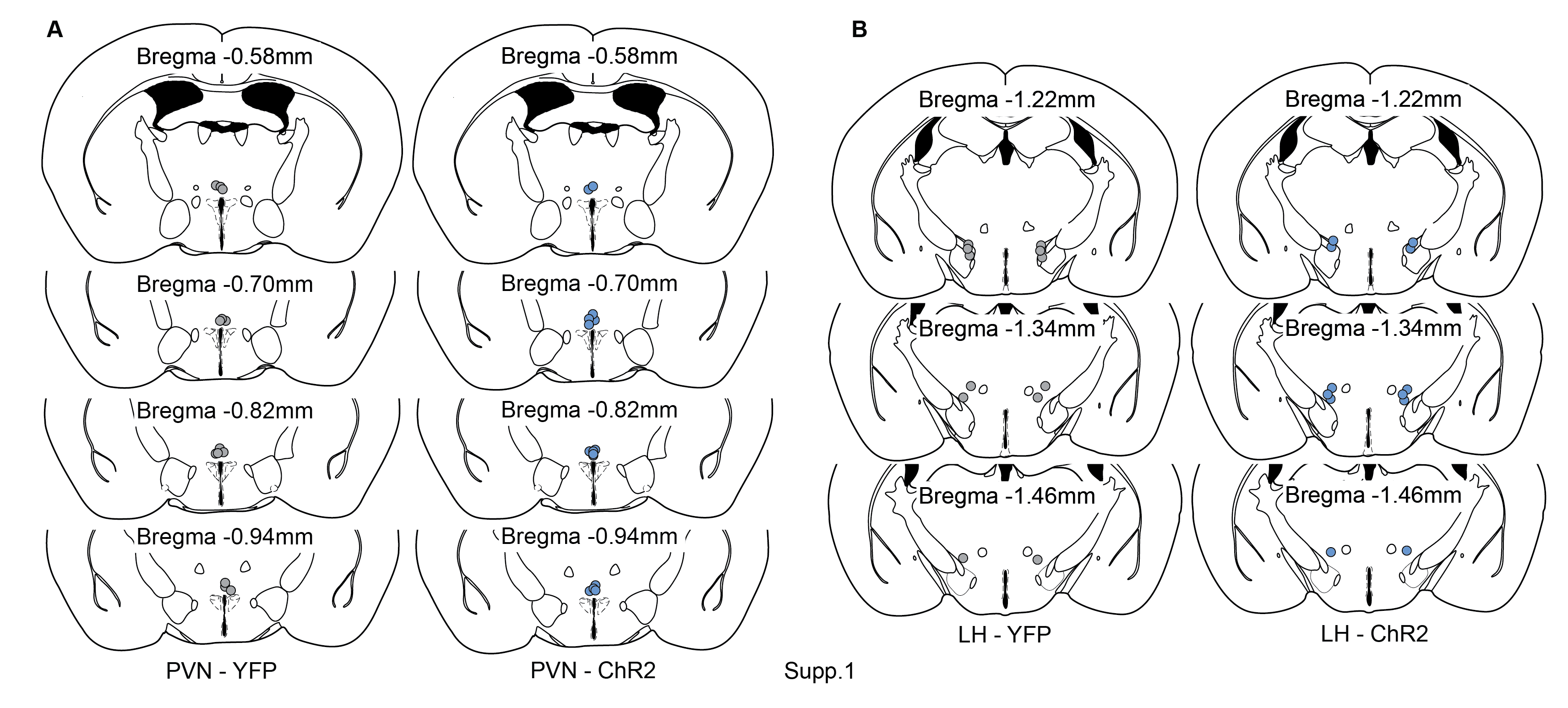

### Supplement Figure 1

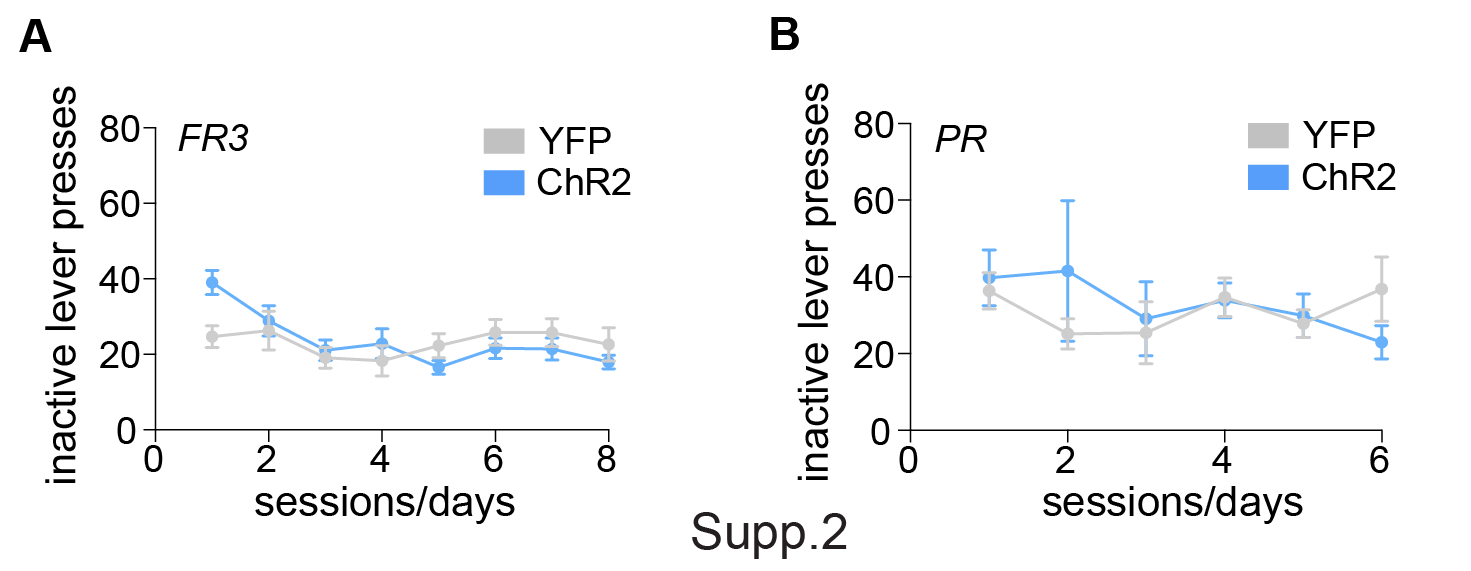
